# 11β-HSD2 buffers fetal glucocorticoid exposure inducing *Per1* expression under maternal stress

**DOI:** 10.64898/2026.04.13.718085

**Authors:** Kazuya Yabumoto, Yasuhiro Umemura, Hitomi Watanabe, Yasuhiro Endo, Nobuya Koike, Akiyo Kakibuchi, Akira Sugimoto, Taisuke Mori, Gen Kondoh, Kazuhiro Yagita

**Affiliations:** Department of Physiology and Systems Bioscience, Kyoto Prefectural University of Medicine, Kyoto 602-8566, Japan; Department of Obstetrics and Gynecology, Graduate School of Medical Science, Kyoto Prefectural University of Medicine, Kyoto 602-8566, Japan; Laboratory of Integrative Biological Science, Institute for Life and Medical Sciences, Kyoto University, Kyoto 606-8507, Japan

**Keywords:** circadian rhythm, maternal–fetal communication, glucocorticoid, 11β-HSD2, segmentation clock

## Abstract

Glucocorticoids (GCs) have been proposed as maternal-fetal communication signals. However, fetal circadian rhythms are initially shielded from maternal entrainment, in addition to delayed circadian clock emergence due to CLOCK suppression. Premature CLOCK/BMAL1 activation disrupts *Hes7*-driven somite-like structure in gastruloids. Given the genomic proximity of *Per1* to *Hes7* and their transcriptional ripple effect, the physiological significance of delayed cell-autonomous circadian clock development and the temporal program of maternal-fetal communication during the developmental process have remained unclear. Here, based on a marked decline in *Hsd11b2*, encoding a GC-inactivating 11β-HSD2 enzyme, during organogenesis, we performed split-litter embryo-transfer experiments in which *Hsd11b2* knockout (KO) and wild-type (WT) embryos shared the same maternal environment. Amniotic fluid (AF) GCs remained low and arrhythmic under basal conditions. In contrast, maternal stress caused a pronounced GC surge and *Per1* induction in KO, suggesting that 11β-HSD2 buffers acute maternal GC surges. Despite the genomic proximity of *Per1* to *Hes7* and their transcriptional ripple effect, stress-associated and pharmacological GC exposure recapitulated no overt segmentation defects *in vivo*. Embryonic stem cell-derived gastruloid assays confirmed that neither GC exposure nor *Per1* induction arrested *Hes7* oscillations, whereas premature CLOCK/BMAL1 activation impaired these processes even in *Hes7* KO gastruloid with ectopic rescue, suggesting that interference with the segmentation clock is mediated by premature CLOCK/BMAL1 activation, not by GC-induced *Per1* expression. These findings clearly show that maternal GC signals are selectively buffered during early development. In addition, suppression of CLOCK/BMAL1 activity preserves segmentation clock function, indicating delayed circadian clock emergence is actively regulated during embryogenesis.

**Significance Statement:** GCs have been proposed as maternal-fetal communication signals. However, initially, circadian clock is not only suppressed but also shielded from maternal entrainment. Premature CLOCK/BMAL1 activation can disrupt *Hes7*-driven somitogenesis. In a split-litter *Hsd11b2*-knockout model, AF GCs remained low and arrhythmic basally but surged after maternal stress in KO embryos, inducing *Per1*. Despite a genomic position effect of *Per1*–*Hes7* and their putative transcriptional coupling, stress-associated or pharmacological GC exposure did not cause segmentation defects *in vivo* or disrupt *Hes7* oscillations *in vitro*, whereas CLOCK/BMAL1-driven arrest of *Hes7* oscillations persisted in gastruloids despite ectopic *Hes7* rescue. These findings identify 11β-HSD2 as a developmental buffer and support the physiological importance of the temporal architecture controlling the timing of circadian clock development.

## Introduction

Mammalian development is governed not only by spatial patterning that shapes morphology, but also by temporal programs that dictate the timing and sequence of major physiological events(1). Among the temporal programs, the process of mammalian circadian molecular clock establishment during ontogenesis also proceeds along a strictly controlled and species-specific timeline(2). The circadian clock with an approximately 24-h cycle is encoded by transcriptional-translational feedback loops (TTFLs) of core clock genes(3) and oscillates at the cellular level(4–7), playing anticipatory roles in maintaining biological homeostasis(8, 9). The development of the circadian clock has been demonstrated to be based on cell-autonomous mechanisms coupled to cellular differentiation, as shown in the *in vitro* differentiation system using embryonic stem cells (ESCs) and induced pluripotent stem cells (iPSCs) (10). *In vitro* differentiation of mouse ESCs and iPSCs indicates the delayed emergence of the circadian oscillations in culture, even using human iPSCs(11, 12). *In vivo*, the fetal clock begins to oscillate only in the latter half of organogenesis(2, 11), and this delayed onset is similar to that observed in *in vitro* differentiation systems. Furthermore, we reported the possibility that ectopic CLOCK/BMAL1 expression in the early developmental stage with suppressed circadian oscillations interferes with the segmentation clock oscillation, potentially through perturbation of *Hes7*-related signaling pathways (WNT/MAPK/NOTCH) and/or the gene regulation across the mammalian-conserved syntenic region surrounding the *Hes7* locus, including the adjacent core circadian clock gene *Per1*(13). These observations suggest that suppression of circadian clock during early development is an actively regulated and functionally significant process.

On the other hand, the maternal environment exerts a significant impact on mammalian development. At later developmental stages, the fetal circadian clock can be entrained by maternal rhythm(11, 14, 15). Although it has often been assumed that circadian rhythm entrainment between mother and fetus operates throughout pregnancy(16–19), the transcriptional rhythms in the embryonic hearts at E10–E12 are hardly detectable, suggesting that embryonic tissues during early development may be shielded from maternal entrainment signals(11). However, the mechanisms governing the establishment of circadian maternal–fetal entrainment remain unknown.

Glucocorticoids (GCs) may contribute to maternal–fetal communication(20) and stand out among candidate entrainment signals such as humoral factors or body temperature cycles(21). Although maternal GC dynamics are strictly species-dependent, a marked increase in GC levels after organogenesis is conserved in both humans and mice(22, 23).

Before this, fetal adrenal GC production is immature(24, 25), making maternal placental transfer the main source of fetal exposure. The maternal GCs may mediate prenatal stress effects(26–29), with some studies reporting skeletal patterning defects by unknown mechanisms(30). Meanwhile, because GC signaling can also rapidly induce expression of circadian genes including *Per1* (31, 32), stress-associated maternal GC elevations during pregnancy could influence the embryonic circadian clock. However, GC dynamics within embryos during organogenesis, particularly under basal and stress conditions, and their impact on embryonic circadian and developmental programs remain unclear.

One of the key factors limiting the maternal–fetal GC transfer is 11β-hydroxysteroid dehydrogenase type 2 (11β-HSD2; encoded by *Hsd11b2* gene), which inactivates GCs(33, 34). *Hsd11b2* is expressed during organogenesis in both humans and mice. In humans, 11β-HSD2 appears in placental syncytiotrophoblasts around three weeks after fertilization(35), with expression and activity fluctuating during pregnancy(36–38). In mice, *Hsd11b2* mRNA is widely expressed in the embryo, including the placenta, but wanes within a few days after organogenesis(39). Although post-organogenesis expression patterns differ between mice and humans(35, 39), *Hsd11b2* expression or 11β-HSD2 activity declines near term in both species(38, 39), implying that fetal GC exposure is consistently regulated by developmental stage and gestational timing.

In the present study, upon reanalyzing our previously published data(11), *Hsd11b2* showed a marked change in murine embryonic hearts during organogenesis. Based on this finding, using a split-litter embryo-transfer experimental design comparing *Hsd11b2* knockout (KO) and wild-type (WT) embryos, we rigorously profiled maternal–fetal GC dynamics and embryonic transcriptional responses during organogenesis under basal conditions and after acute maternal restraint stress. Under the basal conditions, amniotic fluid (AF) GCs remained low and arrhythmic in both genotypes, despite the robust maternal circadian rhythm. Under the stress conditions, the AF GC levels rose markedly in the KO embryos, suggesting that while multiple mechanisms not limited to the 11β-HSD2 function may act to suppress GC exposure to the embryo under basal conditions, 11β-HSD2 is essential as a surge buffer under maternal acute stress. Furthermore, surprisingly, the maternal stress was associated with *Per1* induction in KO embryonic hearts *in vivo*, prompting further investigation into whether the stress-associated *Per1* response might affect fetal morphogenesis, potentially by a genomic position effect of *Per1*–*Hes7*(13, 40). However, across all stress conditions tested, we did not observe overt segmentation defects in the split-litter model. In addition, ESC-derived gastruloid assays also showed that neither GC exposure nor *Per1* induction affected autonomous *Hes7* oscillations, whereas forced CLOCK/BMAL1 expression impaired them even under ectopic *Hes7* KO-rescue conditions. These results suggest that the suppression of functional CLOCK/BMAL1 during early development may be important for proper somitogenesis. Together, these findings provide insight into how maternal signals are selectively gated during development and support the idea that delayed emergence of circadian oscillations is an actively maintained feature of embryogenesis.

## Results

### CRISPR/Cas9 excision yields an *Hsd11b2* knockout mouse model

Firstly, we evaluated embryonic stage–dependent gene expression changes in the murine embryonic hearts by reanalyzing our previously published dataset(11) and then found that *Hsd11b2* was ranked highly in ascending order of gene expression change (Fig. 1A, Fig. S1, Dataset S1). The *Hsd11b2* expression in the embryonic hearts was decreased along the course of organogenesis at E10–E12 and declined significantly at E17–E19 (Fig. 1B). We then generated and validated an *Hsd11b2*-knockout mouse line. Using the CRISPR/Cas9 system, an ∼4.2-kb deletion spanning exons 1–5 was introduced into the highly prolific outbred ICR strain and verified by Sanger sequencing (Fig. 1C, D). Duplex PCR assay clearly separated WT, heterozygous (Het), and homozygous (KO) alleles (Fig. 1E), and the genotype frequencies followed Mendelian expectations (χ^2^ = 4.66, df = 2, *P* = 0.10). The immunoblotting of embryonic heart extracts confirmed that the 11β-HSD2 protein was undetectable in the homozygous knockouts, whereas the WT littermates showed the expected ∼40-kDa band (Fig. 1F). These data demonstrate the establishment of an *Hsd11b2* knockout line for downstream phenotypic analyses.

**Figure 1.**
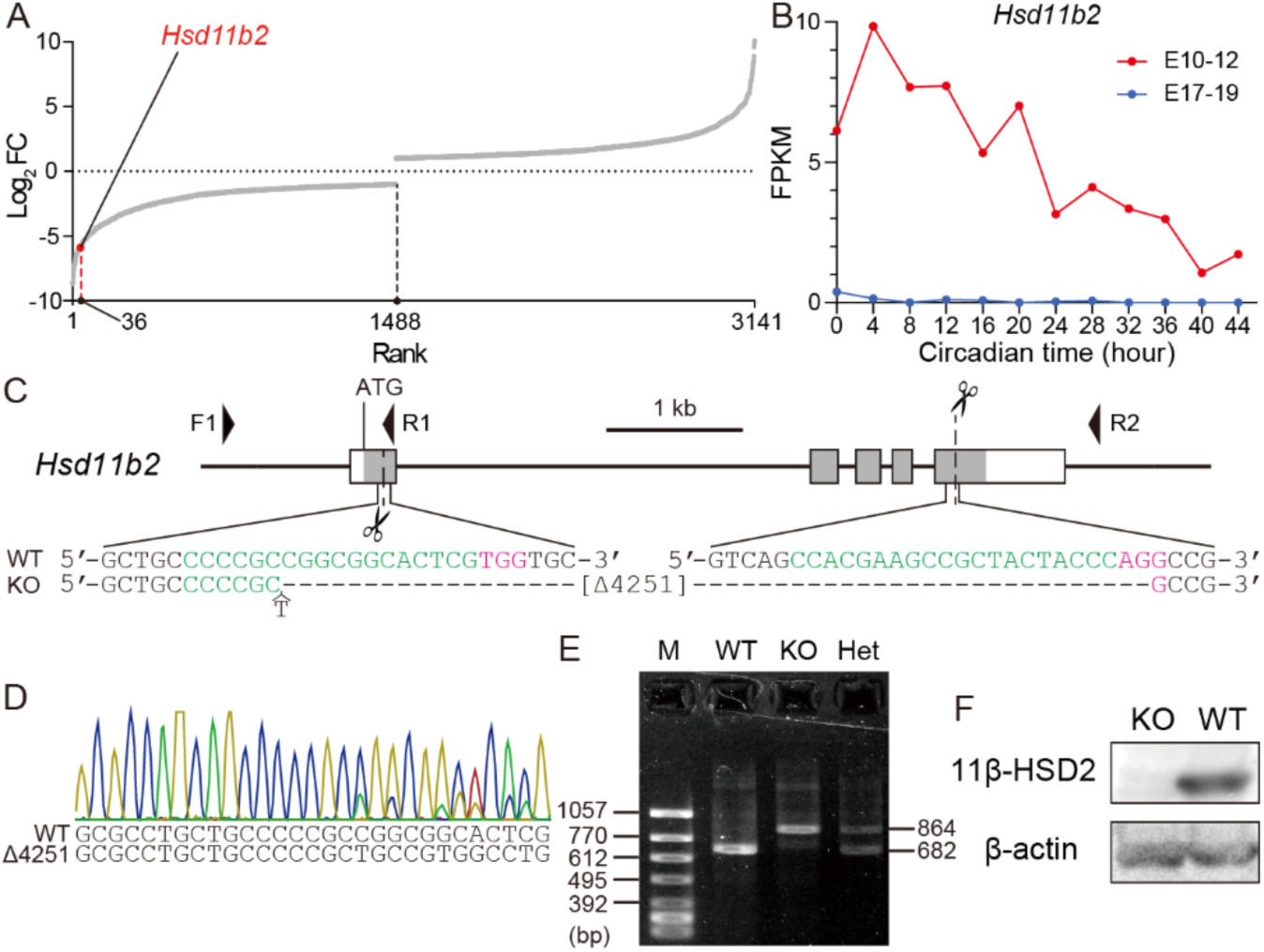
Generation and validation of the *Hsd11b2* KO mouse line. (A) Rank plot of log_2_ fold change (E10–12 vs. E17–19) reanalyzed from RNA-seq data of embryonic hearts (DRA005754) (11). Only genes with absolute fold change > 2 and FDR < 0.05 are indicated. Negative values indicate lower gene expression at E17–19 relative to E10–12. (B) Time-course expression of *Hsd11b2* (DRA005754)(11). (C) Schematic representation of the *Hsd11b2* target regions and DNA sequences of wild type (WT) and knockout (KO) mice. The boxes represent exons and the coding region marked in gray. The target DNA sequences for guide RNAs and protospacer adjacent motif (PAM) are indicated in green and red, respectively. The dashed line denotes the deleted region (Δ4,251 bp). Arrowheads indicate the positions of the genotyping primers (not drawn to scale). The horizontal scale bar represents 1 kb. (D) Sanger sequencing confirmed the precise 4,251-bp deletion. The chromatogram illustrates the cut sites in a heterozygous *Hsd11b2^+/–^*founder. (E) Allele-specific PCR for routine genotyping. Primer set F1/R1 amplifies a 682-bp wild-type fragment; F1/R2 amplifies an 864-bp knockout fragment. Representative agarose gel electrophoresis image shows DNA ladder (M), WT (+/+), homozygous knockout (KO, −/−), and heterozygote (Het, +/−). (F) Western blot of E10.5 embryonic heart lysates demonstrates loss of 11β-HSD2 protein in *Hsd11b2* KO embryos.

### Embryonic AF GC is low and arrhythmic; 11β-HSD2 loss produces only a modest baseline increase

To determine whether maternal GCs and the fetal and placental enzyme 11β-HSD2 participate in circadian maternal–fetal entrainment, we profiled GC dynamics and embryonic cardiac gene expression in a 4-h time series. WT and KO zygotes were separately implanted into the opposite oviducts of the same pseudopregnant dam, creating a split-litter embryo-transfer experimental design that exposed both genotypes to an identical maternal environment (Fig. 2A). As a validation of our experimental system in the outbred ICR background, we confirmed the expected gestational increase of corticosterone levels in maternal serum and AF (Fig. 2B). Starting at ZT1 on E10, the maternal serum, AF, maternal and embryonic hearts, and residual embryonic tissues were collected every 4 h for 52 h (Fig. 2C), for hormone assays and RNA-seq analysis. Genotyping of the residual embryonic tissues confirmed that embryos of each genotype were predominantly recovered from the uterine horn ipsilateral to the transfer side. In addition, embryo recovery rates at E10–E12 did not differ between genotypes, indicating that *Hsd11b2* deficiency does not cause overt embryonic lethality during organogenesis (Fig. 2D).

**Figure 2.**
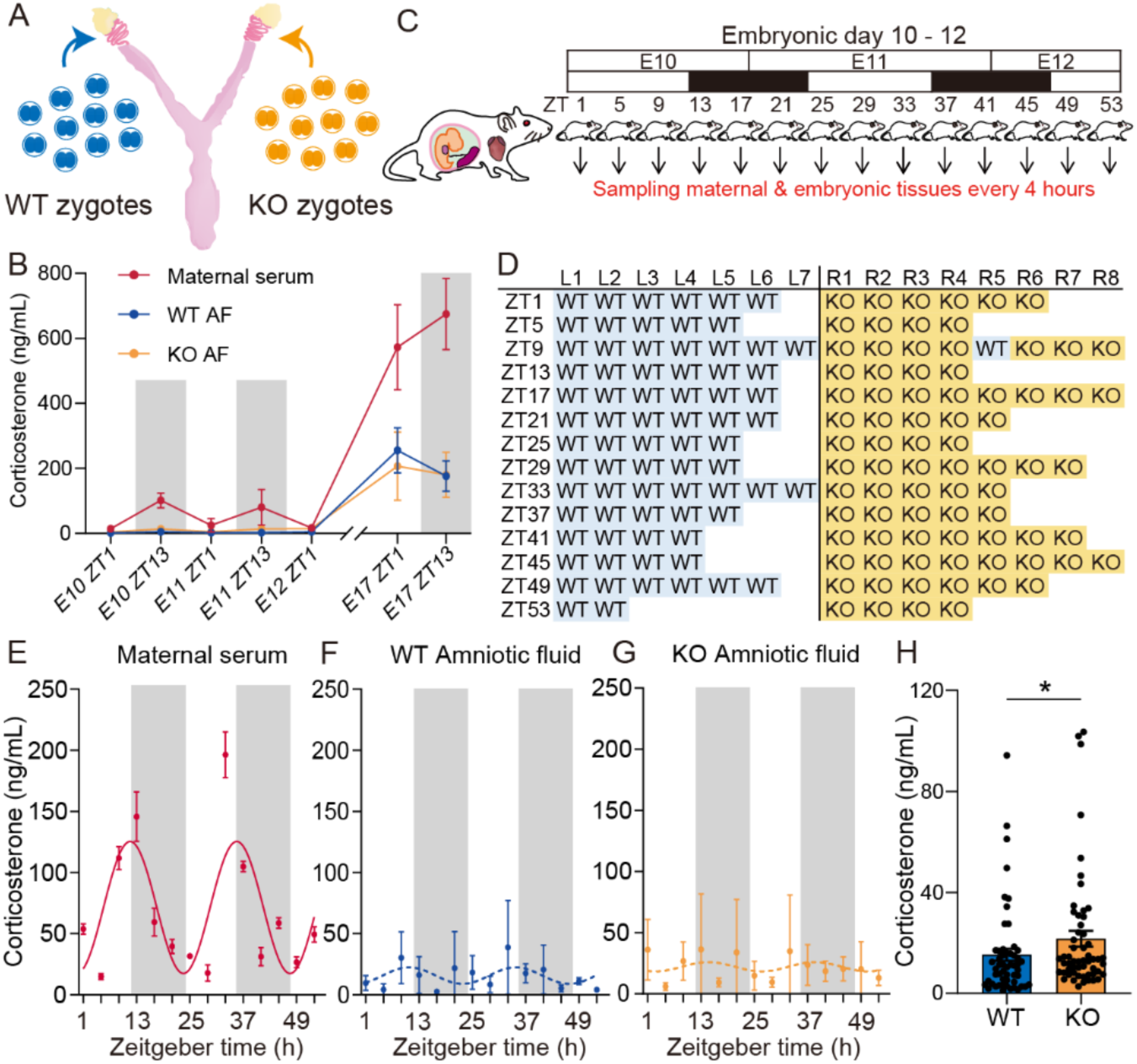
Experimental design of time-course analysis and corticosterone dynamics during gestation. (A) Schematic diagram of the embryo-transfer strategy. WT (blue) and *Hsd11b2* KO (orange) zygotes (≤ 10 per genotype) were transferred into the opposite oviduct of the same pseudopregnant dam, thereby exposing both embryonic genotypes to the same maternal environment. (B) Corticosterone concentrations in the maternal serum and AF at ZT1 and ZT13 in the outbred ICR background. Each point shows mean ± SD (maternal serum, n = 3 dams; AF, n = 9 embryos per genotype). Gray shading indicates the dark phase. (C) Sampling schedule for the time-series experiment. Maternal blood, AF, maternal hearts, embryonic hearts and the residual embryonic tissues were collected every 4 h from ZT1 to ZT53 (E10–E12). White and black bars denote light and dark phases, respectively. (D) Genotyping of embryos collected in the E10–E12 time-series experiment, showing the genotype of each recovered embryo by uterine horn (transfer side) and Zeitgeber time. (E–G) Corticosterone profiles are shown at 4-h intervals. Maternal serum (red) shows a robust circadian rhythm (cosinor analysis, P < 0.0001), whereas AF remains low and arrhythmic in both WT (blue) and KO (orange) embryos (cosinor analysis, P = 0.1505 for WT, P = 0.6919 for KO). At each timepoint, the maternal value was derived from one dam assayed in technical triplicate (mean ± SD), while the embryonic values represented the mean ± SD of 2–4 individual embryos per genotype. Gray shading marks the dark phase. (H) AF corticosterone pooled across all time points. Bars show mean ± SD (WT n = 54, KO n = 56, two-sided Wilcoxon rank-sum test, \**P* < 0.05).

As an initial readout, the corticosterone levels were measured on both sides of the maternal–embryonic interface to determine whether the maternal rhythm entrains the embryonic profile and whether this process is altered by loss of 11β-HSD2. The maternal serum corticosterone displayed a robust 24-h rhythm, peaking shortly after lights-off and troughing at mid-light phase (Fig. 2E), consistent with previous reports in pregnant mice(22). In contrast, the AF corticosterone remained markedly lower than maternal levels throughout the time course and showed no discernible circadian rhythm in either genotype: WT (Fig. 2F) and KO (Fig. 2G). Nevertheless, the AF corticosterone concentrations were slightly but significantly higher in KO than in WT littermates (Fig. 2H). Thus, the loss of 11β-HSD2 induces only a modest increase in embryonic GC exposure in mid-organogenesis, but the robust maternal GC rhythm did not produce a detectable GC rhythm in AF, even in KO embryos.

### Baseline cardiac transcriptome is largely unaffected by 11β-HSD2 loss

Next, to test whether *Hsd11b2* deficiency alters gene expression, we performed RNA-seq analysis of the hearts. Cardiomyocyte markers such as *Mef2c*, *Nkx2-5*, and *Tbx5* were expressed in maternal hearts and embryonic hearts of both genotypes (Fig. S2A). The transcriptional landscapes of WT and KO embryos overlapped extensively: 13,090 expressed genes were detected in both genotypes (Fig. 3A). The UMAP embedding showed no genotype-specific clustering (Fig. 3B). The analysis of differentially expressed genes (DEGs) between WT and KO embryonic hearts identified only 0.1% (11 genes) of the transcriptome (Fig. 3C). The expression of core clock genes was rhythmic in maternal hearts (except *Clock*), whereas none were rhythmic in embryonic hearts of either genotype (Fig. S3). The *Hsd11b2* mRNA was virtually absent in the KO hearts, and it declined over time in the WT hearts, similar to the pattern observed in the C57BL/6J strain (Fig. 3D, Fig. 1B). To exclude the possibility that this minimal change in the transcriptome reflected an absence of GR (*Nr3c1*), we examined its expression: *Nr3c1* was expressed in both genotypes (Fig. 3E), and the GR protein was readily detected by western blot in hearts of both WT and KO embryos (Fig. 3F). These results indicate that, under normal maternal conditions, *Hsd11b2* deficiency alone does not appreciably disrupt the embryonic cardiac gene expression network during organogenesis.

**Figure 3.**
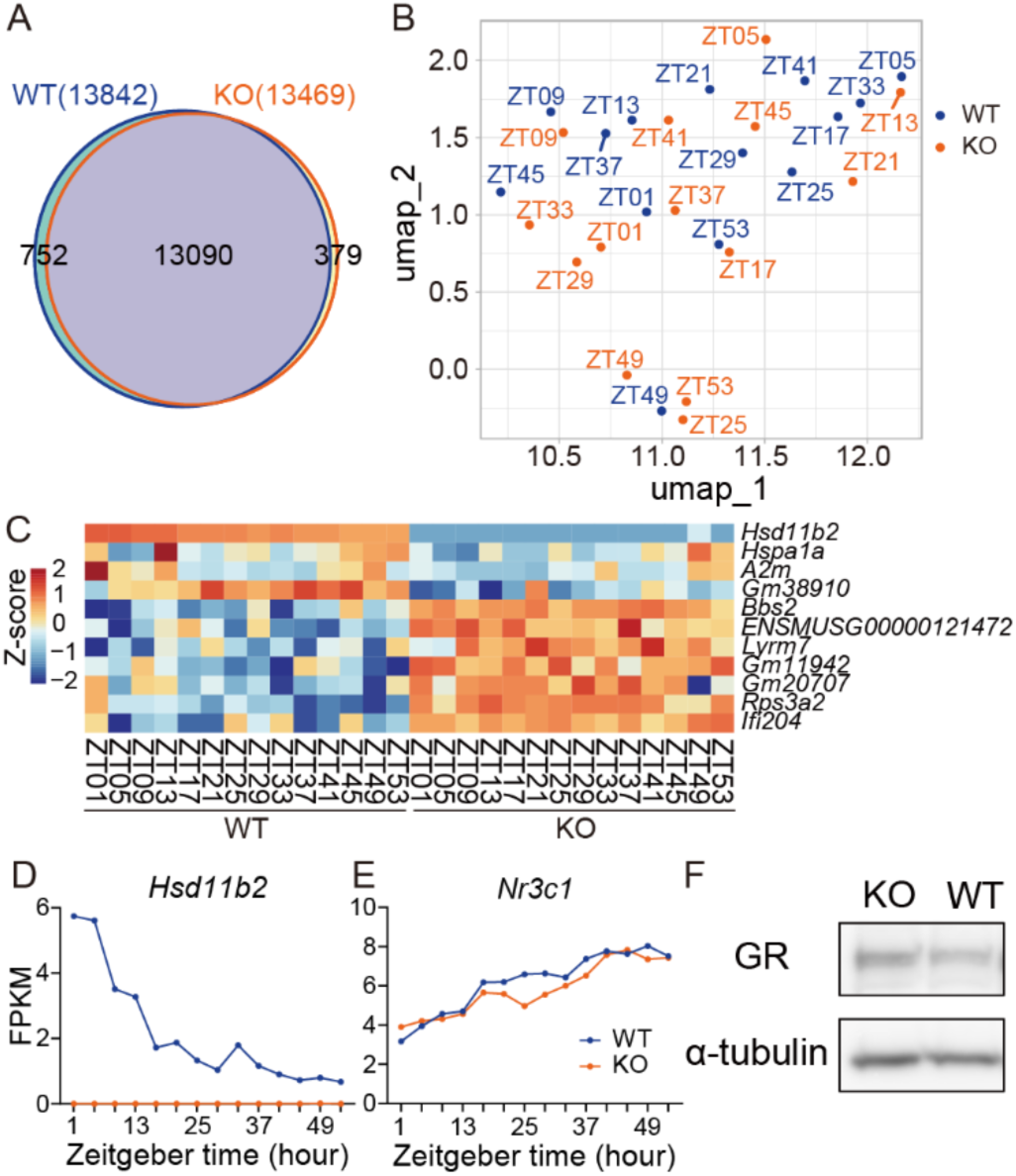
Transcriptomic analysis of embryonic hearts at E10–12. (A) Venn diagram of genes expressed (FPKM > 0.5; ≥ 20 raw reads) in WT and KO embryonic hearts. (B) UMAP of embryonic heart samples colored by genotype (WT, blue; KO, orange). (C) Heatmap view of DEGs between WT and KO embryonic hearts (FDR < 0.05); rows represent genes, columns represent individual samples, and the color scale denotes z-score. (D, E) Temporal expression profiles of *Hsd11b2* and *Nr3c1* in embryonic hearts (E10–E12) plotted aligned to Zeitgeber time (ZT1–ZT53; 4-h intervals). (F) Western blot of embryonic heart lysates confirmed glucocorticoid receptor (GR) proteins in both genotypes.

### Acute maternal stress unmasks a strong genotype × stress interaction in AF corticosterone

Because *Hsd11b2* deficiency caused only a slight increase in AF corticosterone and virtually no transcriptional alterations under basal maternal conditions, we next investigated whether 11β-HSD2 would become indispensable when maternal GCs surge acutely. Using the same split-litter experimental design described above (Fig. 2A), WT and KO zygotes were placed separately in the opposite oviducts of a single dam, and the pregnant females were either left undisturbed or subjected to a 2-h restraint on E11.5 (Fig. 4A).

**Figure 4.**
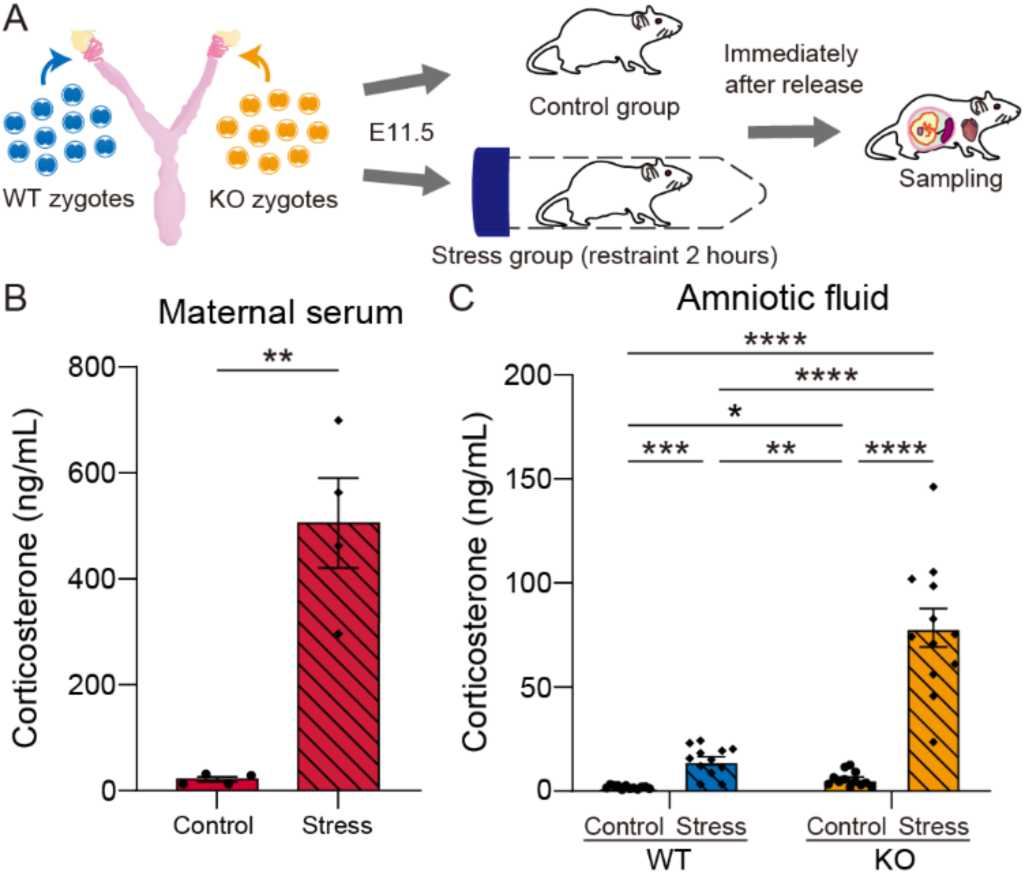
Acute maternal restraint stress and corticosterone levels. (A) Experimental outline. Dams carrying the embryo-transfer litters described in Fig. 2A were either left undisturbed (control) or restrained for 2 h (stress) at E11.5. (B) Maternal serum corticosterone was measured immediately after restraint. The solid bar indicates the control group and the hatched bar shows the stress group. Bars show mean ± SD (n = 4 dams per group, two-sided Welch’s *t* test, **P < 0.01). (C) AF corticosterone in WT and KO embryos under control (solid bars) and stress (hatched bars) conditions. Bars show mean ± SD (n = 12 embryos per group). Two-way Welch–James heteroscedastic ANOVA revealed main effects of genotype (F(1, 12.35) = 50.50, P < 0.0001) and stress (F(1, 12.35) = 80.14, P < 0.0001), and significant interaction between them (F(1, 12.35) = 40.29, P < 0.0001). Games–Howell post-hoc tests: *P < 0.05, **P < 0.01, ***P < 0.001, ****P < 0.0001).

The restraint provoked the expected maternal endocrine response and the serum corticosterone increased by almost 20-fold relative to control conditions (Fig. 4B). In AF, a pronounced genotype × treatment interaction emerged. Under the basal conditions, KO embryos displayed a small but significant elevation over WT littermates. The acute stress raised AF corticosterone about 6-fold in WT embryos, yet produced a striking ∼14-fold rise in KO embryos (Fig. 4C). Two-way Welch–James ANOVA confirmed the significant main effects of genotype and stress and a highly significant interaction. Thus, although 11β-HSD2 deficiency causes only a modest baseline increase of AF GC under basal maternal conditions, it allows a disproportionately large maternal GC pulse to reach the embryo during maternal acute stress, suggesting that 11β-HSD2 functions as a critical barrier against transient maternal GC surges in mid-organogenesis.

### Loss of 11β-HSD2 broadens embryonic cardiac transcriptional responses to acute maternal stress

Next, we profiled genome-wide transcriptional changes immediately after 2 h of acute restraint stress by RNA-seq in both dams and embryos. After the cardiomyocyte markers were verified (Fig. S2B), the 311 DEGs were detected in the maternal hearts, and the upregulated set included classical primary GC target genes such as *Fkbp5*(41), *Tsc22d3*(42), and *Per1*(32) (Fig. 5A and B, Dataset S2). These results confirmed that GC signaling was activated in the maternal compartment during the restraint stress.

**Figure 5.**
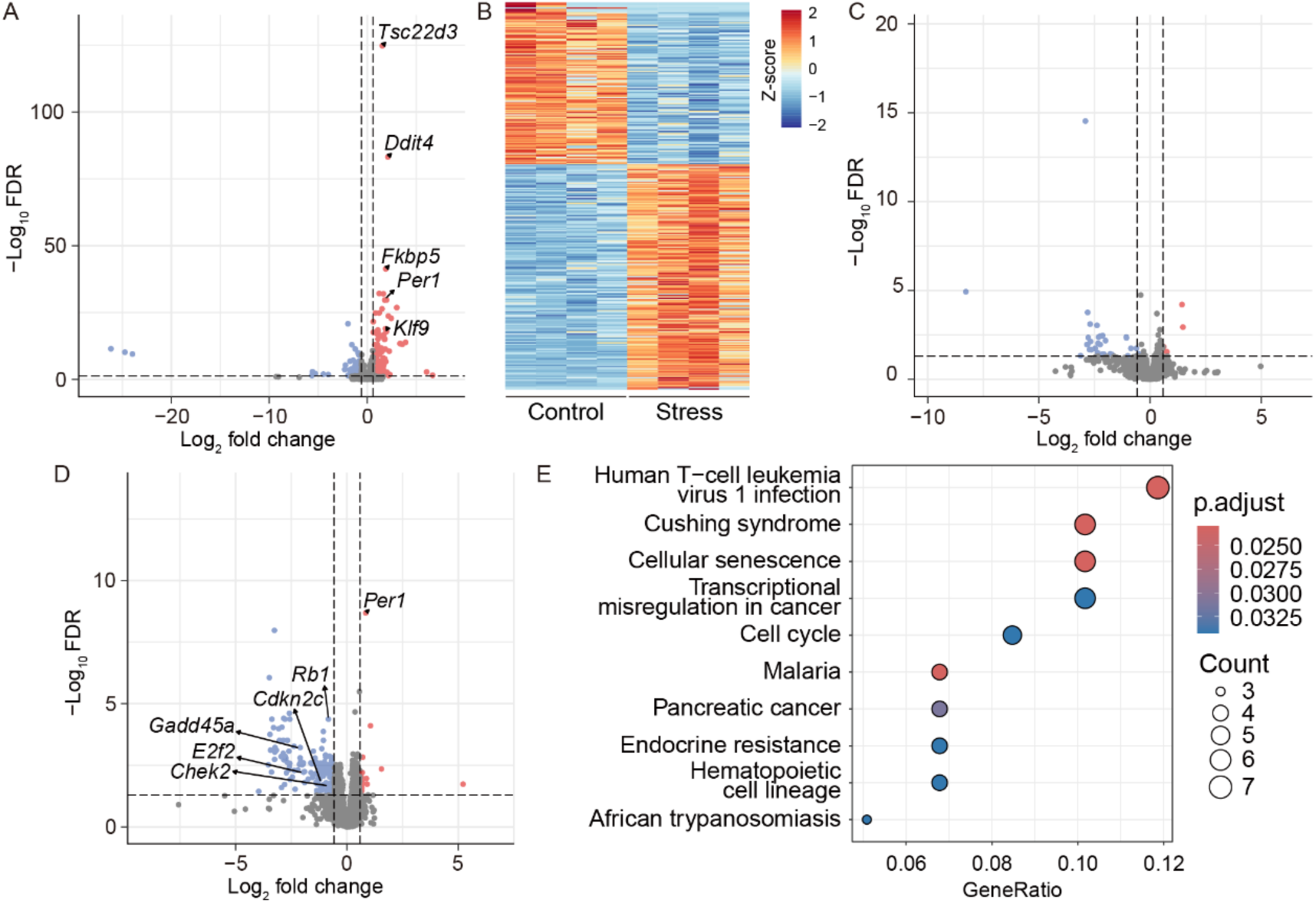
Transcriptomic responses to acute maternal stress at E11.5. (A-D) Volcano plots and the heatmap view of stress-induced differentially expressed genes (Stress/Control) in (A, B) maternal hearts, (C) WT embryonic hearts, and (D) KO embryonic hearts after 2-h restraint stress. The x-axis indicates log_2_ fold change (Stress/Control) and the y-axis indicates −log_10_(FDR). The horizontal dashed line marks FDR = 0.05; vertical dashed lines denote |log₂FC| = 0.585 (FC = 1.5). DEGs were defined with DESeq2 at FDR < 0.05 and absolute fold-change > 1.5. (E) Dot plot of top 10 enriched KEGG pathways among the DEGs downregulated in *Hsd11b2*^−/−^ embryonic hearts under stress.

In contrast, the embryonic cardiac responses differed between the genotypes. In the WT embryos, the stress-induced transcriptional changes remained relatively limited with only 37 DEGs and no canonical GC-responsive genes were detected among the upregulated transcripts (Fig. 5C). In the KO, however, 169 DEGs were detected, highlighting a markedly broader transcriptional response than in the WT, including the significant induction of primary GC target *Per1* (Fig. 5D). KEGG pathway analysis showed significant enrichment of cell-cycle pathways among genes downregulated in the KO hearts under the stress, including Rb-E2F axis genes (*Rb1*, *E2f2*) and additional cell-cycle regulators (Fig. 5E, Dataset S3). In the comparison between stressed WT and KO embryonic hearts, only 6 genes were identified as DEGs; among these, *Per1* showed significantly higher expression in KO hearts (Fig. S4). Collectively, these results suggest that, under the acute stress, the loss of 11β-HSD2 permits excessive maternal GC surges to reach the embryos, leading to the pronounced transcriptional alterations that encompass the induction of GC target genes and the repression of cell-cycle regulators in the embryonic heart.

### CLOCK/BMAL1-driven disruption of somitogenesis-like processes in gastruloids does not require *Per1*–*Hes7* genomic adjacency

Given that *Per1* is genomically adjacent to the segmentation clock gene *Hes7*, we further investigated whether stress-induced *Per1* activation perturbs somitogenesis via transcriptional ripple effects(13, 40). In the split-litter model, we subjected pregnant dams to maternal restraint stress or corticosterone administration from E9.5 to E12.5 and assessed E14.5 fetuses by cartilage staining, focusing on the vertebral column. However, across all conditions tested, we did not observe overt segmentation defects (Fig. S5).

The stress-associated *Per1* induction detected in embryonic hearts was not examined in somitogenesis-relevant tissues including presomitic mesoderm. We next assessed the effects of GC-related perturbations on somitogenesis *in vitro* using gastruloids, ESC-derived embryonic organoids that recapitulate a somitogenesis-like process(43). Dexamethasone (Dex) stimulation, largely resistant to 11β-HSD2 activity, failed to perturb the autonomous *Hes7* oscillations (Fig. 6A–C). The Dex stimulation in the gastruloids did not significantly induce *Per1* mRNA despite low but detectable *Nr3c1* expression (Fig. 6D, Fig. S6A). Because the lack of *Per1* induction upon Dex treatment may reflect insufficient activation of GR signaling at the developmental stage represented by gastruloids, we performed forskolin (Fsk) stimulations in the gastruloids. *Per1* is known to be acutely induced by Fsk via cAMP/CREB signaling(44). Fsk stimulation in gastruloids induced *Per1* expression (Fig. 6E), but also failed to perturb the autonomous *Hes7* oscillations (Fig. 6F–H).

**Figure 6.**
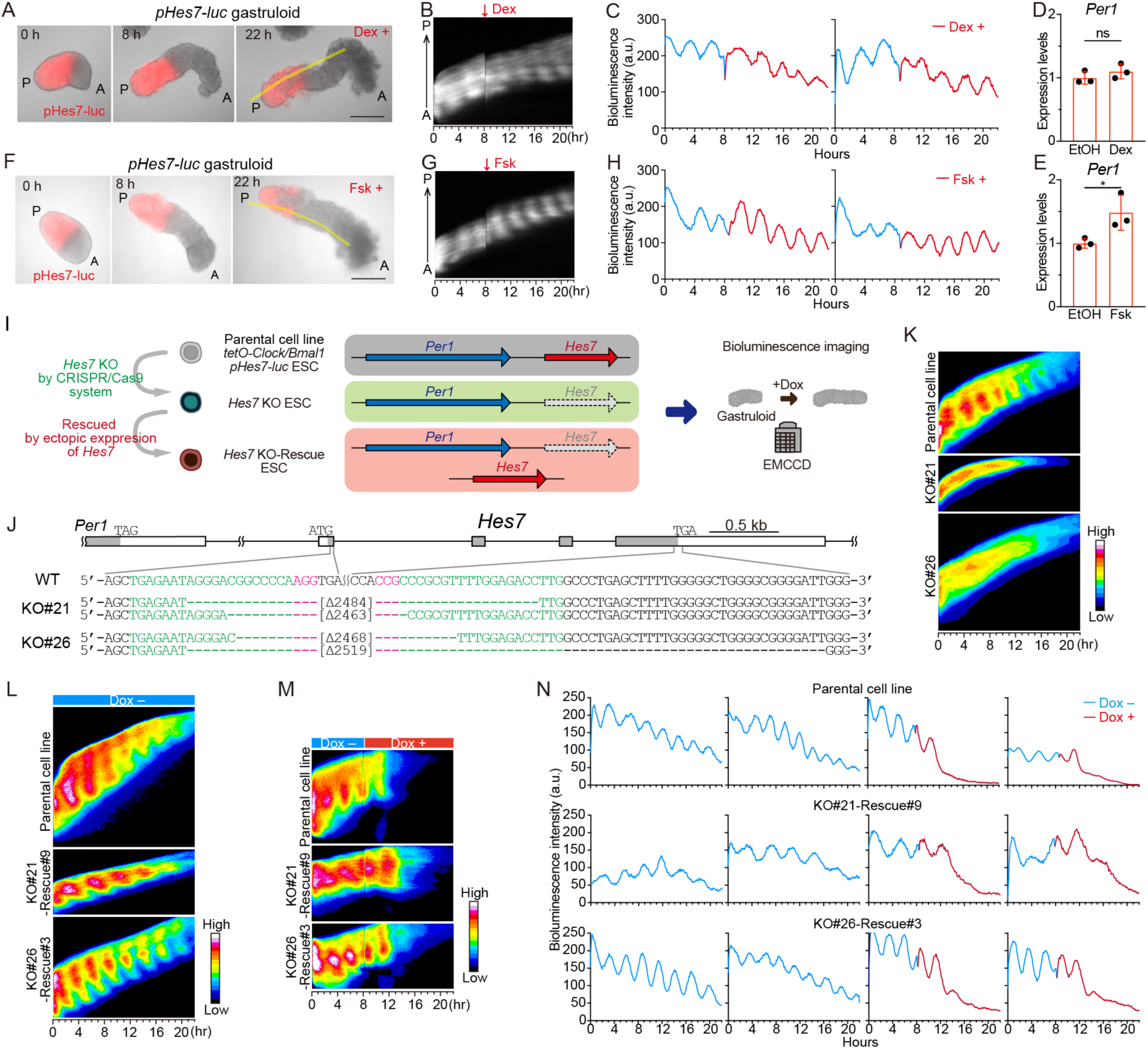
Gastruloid assays of a somitogenesis-like process using Dex/Fsk treatment and *Hes7* KO/rescue lines with inducible CLOCK/BMAL1. (A-C) Time-lapse bioluminescence (red) and bright field imaging of the *pHes7-luc* gastruloid with 300 nM Dex (A). Scale bars = 250 µm. Each kymograph is indicated along the yellow lines in A. Dex-containing medium was added at the indicated time points (B). The representative bioluminescence traces were indicated (C). n = 3 biological replicates. (D, E) qPCR of *Per1* mRNA in the gastruloids after 8 h of culture in 10% Matrigel were treated with 300 nM Dex, 10 µM Fsk, or equal volumes of ethanol (EtOH) to a final 0.1% for 2-3 h. n = 3 biological replicates. Two-tailed t-test. *P < 0.05. ns indicates not significant. (F-H) Same as A–C, but with 10 µM Fsk treatment. (I) *Hes7* KO-rescue cell lines were established using Dox-inducible *Clock/Bmal1 pHes7-luc* ESCs. After the endogenous *Hes7* genes were deleted using the CRISPR/Cas9 system and the *Hes7*-KO ESCs were rescued by the ectopic expression of a *Hes7* promoter-driven *Hes7* gene. The gastruloids derived from these ESC lines were embedded in 10% Matrigel and observed using an EM-CCD camera. (J) Schematic representation of the *Hes7* target regions and DNA sequences of wild type (WT) and knockout (KO) ES cell lines. The boxes represent exons and the coding region marked in gray. The target DNA sequences for guide RNAs and PAM are indicated in green and red, respectively. (K-M) Kymographs in the gastruloids of the indicated ESC lines. (N) Representative bioluminescence traces of the gastruloids. The medium containing Dox at the final concentration of 1000 ng/mL was added on the way of the *Hes7*-reporter oscillations (right). n = 3 biological replicates.

Furthermore, to test whether CLOCK/BMAL1-associated disruption requires the *Hes7* locus adjacent to *Per1* in the genome, *Hes7* KO-rescued ESC lines were established from parental cell lines, doxycycline (Dox)-inducible *Clock/Bmal1* ESCs carrying *pHes7-luc* (Fig. 6I)(13). Almost the entire *Hes7* gene was deleted using CRISPR/Cas9 (Fig. 6J), and then rescued by ectopic *Hes7* expression vector (45). The *Hes7*-KO ESC-derived gastruloids indicated no autonomous *Hes7* oscillations (Fig. 6K), but the gastruloids derived from *Hes7* KO-rescued ESC lines again exhibited the autonomous *Hes7* oscillations (Fig. 6L, Fig. S6B). However, the gastruloids derived from the *Hes7* KO-rescued lines were still found to show the arrest of autonomous *Hes7* oscillations and the up-regulation of *Hes7* expression upon Dox-induced CLOCK/BMAL1 expression (Fig. 6M, N, Fig. S6B). These results suggest that CLOCK/BMAL1-driven disruption cannot be explained solely by *Per1–Hes7* genomic adjacency, and are consistent with the idea that tight suppression of CLOCK/BMAL1 activity is important for normal somitogenesis.

## Discussion

In the present study, we quantified fetal GC dynamics during murine organogenesis and identified 11β-HSD2 as a key buffer that can dampen acute stress–associated GC surges, thereby protecting embryonic cardiac transcriptional responses. Moreover, although *Hsd11b2*-deficient embryos exhibited stress-associated induction of the clock gene *Per1*, *in vivo* and gastruloid-based assays indicate that neither GC exposure nor *Per1* induction recapitulates disruption of somitogenesis-like processes, whereas forced CLOCK/BMAL1 expression disrupts these processes even under ectopic *Hes7* rescue conditions.

Under the non-stressed conditions, the maternal circadian GC rhythm was maintained, whereas the AF GC levels remained low and arrhythmic in both genotypes, with only a negligible increase observed in the KO embryos. These results suggest that multi-layered GC shielding along the maternal–placental–fetal axis—including maternal GC-binding proteins(46) and GC transport processes(47), in addition to placental 11β-HSD2—limits embryonic exposure to GCs during organogenesis under normal conditions.

Consequently, embryos are likely protected not only from elevated GC levels but also from circadian fluctuations in maternal GC. Indeed, the RNA-seq analysis of embryonic hearts under non-stressed conditions showed that only a very small number of genes were differentially expressed between KO and WT, and no notable changes were seen in the overall transcriptional program.

In the acute stress experiments, the GC levels in AF rose only slightly in the WT embryos but spiked sharply in the KO, revealing a clear genotype–stress interaction. In other words, these findings suggest that, during organogenesis, placental-fetal 11β-HSD2 acts as a “surge buffer” protecting embryos from rapid GC surges triggered by acute maternal stress.

By selectively dampening GC surges on the embryo side without attenuating the maternal stress response, 11β-HSD2 may help maintain embryonic hormonal homeostasis, consistent with biphasic developmental effects of GCs—beneficial when appropriately timed, but adverse when premature, excessive, or prolonged(48, 49).

Moreover, the RNA-seq analysis showed that, under maternal stress conditions, KO embryos exhibited more extensive gene expression changes than WT, suggesting that the larger GC surge in AF of KO was reflected at the transcriptional level. Indeed, the KO embryonic hearts displayed a marked increase in *Per1* expression, which is a GC-responsive immediate-early gene. *Per1* is a core circadian clock gene, and *Per1* induction could serve as a key mediator of circadian phase resetting; in the SCN it is rapidly induced by nocturnal light to reset the clock’s phase(50), and in peripheral tissues it is known to respond to GC pulses(32). The stress challenge elicited the striking induction of *Per1* in KO embryos, suggesting that 11β-HSD2 could also function as a “gatekeeper” that regulates the circadian maternal–fetal entrainment.

In the maternal samples, the gene expression changes (particularly involving GC early-response genes) were evident 2 hours after stress onset, consistent with previous reports(51, 52). However, because GC transfer from mother to fetus can be delayed on the order of tens of minutes(53, 54), the transcriptional response in the embryos may not have fully emerged at that 2-hour time point. We have not yet examined the temporal dynamics of other core clock genes or their downstream networks. Therefore, additional time-series analyses will be essential to verify whether GCs can indeed function as an entrainment signal between mother and fetus. Nevertheless, the robust *Per1* induction observed in the present study is consistent with the possibility that stress-associated maternal GC signaling can engage the fetal clock. Although AF corticosterone serves only as a proxy and may not directly reflect GC levels in embryonic tissues or the circulation, the larger rise in AF corticosterone and the broader cardiac transcriptional response in KO embryos in this study support the use of AF corticosterone as an index of embryonic GC exposure.

Maternal stress during pregnancy has been reported to affect fetal growth and morphogenesis depending on the stress paradigm and gestational stage(26, 27), with some studies reporting patterning abnormalities including axial skeletal defects of the vertebrae and ribs(30). Furthermore, the induction of *Per1* expression after maternal stress might contribute to the skeletal patterning defects by perturbing *Hes7* oscillations through the ripple effect within the syntenic region containing *Per1* and *Hes7* (13, 40). We then evaluated whether stress-associated GC exposure could be linked to segmentation abnormalities *in vivo* and *in vitro*. However, neither overt segmentation defects *in vivo* nor the perturbation of autonomous *Hes7* oscillations *in vitro* were observed. While previous studies have indicated an association between maternal stress and developmental abnormalities under specific stress paradigms, our results suggest that fetal GC exposure may be constrained by multi-layered GC shielding and that axial skeletal patterning mechanisms may retain robustness against acute maternal stress, providing important pathophysiological insight into stress-induced developmental abnormalities.

The *Nr3c1* expression was low in the gastruloids, which is consistent with its limited expression in mouse embryos during somitogenesis (Fig. S7)(55). The lack of *Per1* induction upon Dex treatment may reflect the low abundance of *Nr3c1* together with limited GR-mediated responsiveness at this developmental stage. Although the Fsk stimulation induced *Per1* in the gastruloids, it did not affect *Hes7* oscillations, indicating that the level of *Per1* induction achieved here was insufficient to perturb the segmentation clock. By contrast, using gastruloids with Dox-inducible CLOCK/BMAL1 expression, we found that induction of CLOCK/BMAL1 disrupted the autonomous *Hes7* oscillations even in *Hes7*-knockout gastruloids under ectopic *Hes7* rescue conditions. These results suggest that *Per1*–*Hes7* genomic adjacency may not be required for CLOCK/BMAL1-driven disruption of the segmentation clock and raise the possibility that such disruption primarily involves perturbations in signaling networks regulating *Hes7* oscillations(13).

In the KO embryos under the acute stress, we also observed decreased expression of cell cycle–related genes that govern the G1/S transition (e.g., *E2f2* and *Cdkn2c*). GCs have long been known to exert antiproliferative effects(56, 57); for example, in mouse cells *Cyclin D3* mRNA levels drop by about 50% within 2 hours of Dex treatment(58). Considering this prior knowledge alongside the DEGs identified in our study, it appears that the sharp GC surge may have affected cell cycle control in the embryonic hearts. Beyond this short-term analysis after a single acute stress, further studies are needed to determine whether longer-term outcomes—such as sustained suppression of cell proliferation or delayed organogenesis—could result from such GC exposure.

In summary, our findings lead to four key conclusions: i) 11β-HSD2 establishes a developmental barrier that prevents maternal GC signals from reaching the embryo during early pregnancy, ii) maternal stress can override this barrier only in its absence, inducing *Per1* and a GC-responsive transcriptional program, iii) such induction does not compromise somitogenesis, iv) interference with the segmentation clock is mediated by premature activation of CLOCK/BMAL1, not by GC-induced *Per1* expression.

Collectively, our study provides evidence that the delayed emergence of circadian oscillations is an actively regulated and essential feature of embryogenesis. By uncoupling maternal circadian signals from the fetal transcriptional network, 11β-HSD2 safeguards the developmental timing system and ensures robust progression of somitogenesis.

## Materials and Methods

### Animal ethics statement and housing

All animal procedures were approved by both the Animal Experiment Committees from the Institute for Life and Medical Sciences (LiMe), and Kyoto University for mouse generation and surgery, and the Committee of Kyoto Prefectural University of Medicine for subsequent housing and experimentation. All procedures were conducted in accordance with the institutional guidelines and Guidelines for Proper Conduct of Animal Experiments by the Science Council of Japan. Outbred ICR mice (Slc:ICR) were obtained from Japan SLC, Inc. (Shizuoka, Japan) and were maintained under a 12-h light/12-h dark cycle (lights on at 08:00 and off at 20:00), with ad libitum access to food and water.

### Generation of *Hsd11b2* KO mice

The *Hsd11b2* genomic DNA sequences were applied to the CRISPRDirect site (CRISPRdirect (dbcls.jp)(59)) to find the target sequence for the guide RNA in the CRISPR/Cas9 genome editing procedure. Then, guide RNAs for the four target sequences, namely Region 1: CCCCGCCGGCGGCACTCGTGGTG, Region 2: CGCTGGCCTCAATATCGTAGTGG, Region 3: GCCACTCTTGCGTCACTCGAGGG and Region 4: CCACGAAGCCGCTACTACCCAGG were synthesized (FASMAC) and microinjected into fertilized eggs of ICR mice (CLEA Japan) with tracrRNA (FASMAC) and Cas9 protein (Thermo Fisher), respectively. As a result, three founder mice were obtained, and three *Hsd11b2*-knockout mouse lines were established (KO1, KO2 and KO3). In this study, the KO1 line was mainly used.

### Colony expansion

Heterozygous founders (F_0_) were out-crossed once to wild-type ICR mice to (i) increase the number of *Hsd11b2^+/−^* carriers, (ii) minimize somatic mosaicism, and (iii) confirm germ-line transmission of the edited allele. Verified F_1_ heterozygotes were back-crossed to wild-type ICR mice to generate an enlarged F_2_ cohort of carriers.

### Genotyping (all generations and sampled embryos)

Genomic DNA was isolated from tail biopsies at every generation and from residual embryonic tissue collected after each experimental sampling. PCR was performed with allele-specific primers; F1 5′-CCTCATGCAACTTGGGGACT-3′ / R1 5′-GAACTCCGCAGCTCAGACGTA-3′ (WT 682 bp) and F1 / R2 5′-CTTGAGCCCACACATACCCT-3′ (KO 864 bp) using PrimeSTAR Max DNA Polymerase (Takara). Cycling: 98 °C 10 s, 59 °C 5 s, 72 °C 30 s (34 cycles), final extension 72 °C 1 min. Representative amplicons from the F_0_–F_2_ generations were confirmed by Sanger sequencing. To ensure sample identity, we validated genotypes from genomic DNA PCR and RNA-seq; concordant samples were used for downstream analyses.

### Off-target prediction

Potential off-target sites for each sgRNA were identified using Cas-OFFinder(60) v2.5 against the GRCm39/mm39 genome (≤ 2 mismatches, and/or 1-nt DNA/RNA bulge; PAM = NGG). Each retained sequence (≤ 7 per sgRNA) was queried against the NCBI nt database by BLASTN; no retained site overlapped the annotated coding sequences, and only a single partial hit corresponded to an mRNA, indicating a minimal risk of disrupting coding sequences.

### IVF–ET for colony maintenance and generation of pregnant dams for experiments (F₃ onward)

From the F_3_ generation onward, all embryos for colony maintenance were produced exclusively by *in vitro* fertilization-embryo transfer (IVF–ET) between *Hsd11b2^+/−^*parents, whereas wild-type and homozygous knockout embryos were reserved for phenotypic experiments.

Cauda epididymal sperm were collected from male mice of each genotype and incubated in the HTF medium for 90 min to capacitate. Eggs were collected from super-ovulated female mice and fertilized with 5.0 × 10^5^ sperm mL^-1^ in HTF. After 16 h of incubation in the mWM medium, the 2-cell stage embryos were obtained, and then transferred to the oviduct of pseudo-pregnant females. For one recipient, 10 WT embryos were transferred to the right oviduct and 10 KO embryos to the left, respectively.

### Sampling design

General sample handling: All collections were performed under the LD condition (lights on at ZT0). During the dark phase, all procedures up to euthanasia were performed under infrared night-vision goggles (ATN Night Cougar LT); bright light was switched on only after the dam’s eyes had been covered with aluminum foil to prevent photic stimulation. The embryonic hearts were snap-frozen in liquid nitrogen. The maternal hearts were incubated overnight in RNAlater (Thermo Fisher Scientific) at 4 °C, the solution was removed, and the tissues were stored at −80 °C. The serum and AF samples were clarified by centrifugation (5,000 × g, 10 min, 4 °C), and the supernatants were stored at −80 °C until analysis.

Validation (Fig. 2B): Maternal blood and AF were collected at ZT1 and ZT13 on the indicated gestational days (dams; n = 3 per time point and embryos; n = 9 per genotype per time point). Non-stress time-series (E10–E12): The pregnant dams were sampled every 4 h at 14 consecutive Zeitgeber times (ZT1, 5, 9, …, 53) from E10 to E12. At each time point, one dam was removed from the cage and immediately euthanized by cervical dislocation. Maternal blood, AF, maternal and embryonic heart tissues, and residual embryonic tissues for genotyping were collected.

Acute restraint stress experiments: The pregnant dams at E11.5 were restrained for 2 hours starting at ZT23 (1 h before lights-on) in ventilated 50-mL polypropylene tubes. Restraint ended at ZT1, and the samples including maternal hearts were collected immediately after release, following the procedures described above for non-stressed collections.

Repeated maternal treatments for fetal cartilage staining: Pregnant dams were assigned to four groups and treated daily from E9.5 to E12.5 (ZT0 = lights on): (i) stress, 2-h tube restraint at ZT1, ZT4, and ZT7; (ii) control, no restraint or injections; (iii) corticosterone, intraperitoneal corticosterone (20 mg/kg; FUJIFILM Wako Pure Chemical) at ZT1 and ZT9, dissolved in saline containing 10% hydroxypropyl-β-cyclodextrin (HP-β-CD); and (iv) vehicle, injections of 10% HP-β-CD in saline on the same schedule. Embryos were collected at E14.5; the left forelimb was excised for genotyping, and the remaining embryo was fixed and processed for Alcian Blue cartilage staining.

Details of western blotting, corticosterone assay, RNA-seq, cartilage staining of E14.5 embryos, cell culture, transfection, and establishment of Hes7 KO-rescue cell lines, bioluminescence time lapse imaging and data analysis of gastruloid, quantitative RT-PCR (qPCR) and statistical analysis, are included in *SI Appendix*.

## Author Contributions

K.Yagita conceptualized the study; K.Yabumoto, Y.U. and K.Yagita designed the experiments; K.Yabumoto, Y.U., H.W., Y.E., N.K., A.K., A.S. and G.K. conducted the experiments; N.K. analyzed RNA-seq data; T.M. and K.Yagita supervised the study; K.Yabumoto, Y.U., N.K., G.K. and K.Yagita wrote the paper; All authors reviewed and approved the paper.

## Data, Materials, and Software Availability

Data are available in the paper. The raw and processed RNA-seq data reported in this paper are made publicly available via National Center for Biotechnology Information’s Gene Expression Omnibus (GEO) accession number XXXX. Any additional information required to reanalyze the data reported in this paper is available from the corresponding author upon reasonable request.

## Acknowledgments

We thank Yagita lab members for technical assistance and helpful discussion. This work was supported by the Cooperative Research Program (Joint Usage/Research Center program) of Institute for Life and Medical Sciences, Kyoto University, and Grants-in-Aid from the Japan Society for the Promotion of Science (JSPS) to Y.U. (19K06679, 24K09535) and K.Yagita (18H02600, 21H02664, 24H02301).

## Supplementary Information

### SI Materials and Methods

#### Western blotting

Embryonic hearts were homogenized in a buffer containing 20 mM Hepes (pH 7.5), 150 mM NaCl, 10% glycerol, 0.5% Triton-X100, 1 mM DTT, cOmplete Mini protease inhibitor (Roche), 1 mM EDTA, phosphatase inhibitor cocktail (Nacalai Tesque). Lysates were sonicated and clarified by centrifugation. The samples were resolved on SDS/PAGE using 7.5% polyacrylamide gels and were blotted with primary antibodies: anti-11β-HSD2 antibody (1:5,000) (Proteintech), anti-Glucocorticoid receptor (1:5,000) (Proteintech), anti-β-actin (1:25,000) (Sigma-Aldrich), anti-α-tubulin (1:5,000) (MBL).

#### Corticosterone assay

Serum and AF corticosterone were quantified using the Enzo Corticosterone ELISA kit (ADI-900-097). Serum samples were analyzed in technical triplicate and averaged. For AF, 3–4 individual embryos of the same genotype in each litter were assayed once each; these embryonic values were treated as independent biological replicates. Concentrations were interpolated from a four-parameter logistic curve, analyzed with GainData (Arigo Biolaboratories).

#### RNA-seq

Frozen maternal hearts were homogenized three times for 2 min in TRIzol reagent (Thermo Fisher Scientific) with two 7-mm-diameter stainless beads using a desktop bead mill TissueLyser LT (Qiagen) at 50 Hz. Embryonic hearts from 2–4 embryos per litter were pooled for RNA extraction, because total RNA yield from a single embryo was typically ∼600 ng which is insufficient to meet the required input of 1 µg, and to reduce variability arising from dissection and inter-individual differences. Total RNA was extracted using RNeasy columns (Qiagen) according to the manufacturer’s instructions. Poly(A)-enriched, stranded RNA sequencing was carried out by Macrogen Japan Inc. (Tokyo, Japan) on an Illumina NovaSeq X Plus with 101-bp paired-end reads. After adapter sequences were trimmed using Trimmomatic(1), the sequence reads were mapped to the mouse genome (GRCm39/mm39) using STAR(2) as described previously(3). To obtain reliable alignments, the reads with a mapping quality of less than 10 were removed by SAM tools(4). The University of California, Santa Cruz (UCSC) known canonical gene set (57,186) was used for annotation, and the reads mapped to the exons were quantified using the Homer software with an option of -rpkm or -noadj for FPKM or raw read counts, respectively(5). Since the Homer treats each half of the read separately and count each as 0.5 reads for paired-end reads, the raw read counts were rounded to the nearest integer before transforming rlog using DESeq2(6). To report one isoform per locus (gene symbol), the longest expressed isoform was chosen. We assumed that a gene was expressed if there were more than 20 reads mapped on average in the exons of the gene. The expression level cutoff, average FPKM > 0.5, was used for the downstream data analysis. In the embryonic heart dataset from the acute-stress experiment, genes whose maximum Cook’s distance across samples exceeded 0.5 were excluded from downstream analyses(6). Differentially expressed genes in the RNA-seq data were determined using DESeq2 with thresholds of false discovery rate (FDR) < 0.05 and absolute fold change > 1.5. KEGG pathway enrichment analysis of differentially expressed genes was carried out using clusterprofiler in R language(7) with a threshold of false discovery rate (FDR) < 0.05.

#### Cartilage staining of E14.5 embryos

E14.5 embryos were collected and fixed overnight in Bouin’s solution. Samples were then washed several times in 70% ethanol containing 0.1% ammonium hydroxide, equilibrated in 5% acetic acid, and stained with Alcian Blue staining solution (0.05% Alcian Blue in 5% acetic acid) for 2 days. After staining, embryos were rinsed in 5% acetic acid, washed in methanol, cleared in a 1:2 mixture of benzyl alcohol and benzyl benzoate (BABB), and visualized under a stereomicroscope.

#### Cell culture, Transfection, and Establishment of *Hes7* KO-Rescue cell lines

All ESCs were maintained without feeder cells in DMEM (Nacalai) supplemented with 15% fetal bovine serum (Hyclone), 2 mM L-glutamine (Nacalai), 1 mM nonessential amino acids (Nacalai), 100 µM StemSure^®^ 2-mercaptoethanol solution (Wako), 1 mM sodium pyruvate (Nacalai), 100 units/mL penicillin and streptomycin (Nacalai), 1000 units/mL leukemia inhibitory factor (Millipore), 3 µM CHIRON99021 (Tocris Biosciences), and 1 µM PD0325901 (Wako) with 5% CO_2_ at 37°C. *Hes7* KO ESC lines were established using the CRISPR/Cas9 system. E14Tg2a ESCs carrying *Hes7* promoter-driven luciferase reporters and Dox-inducible *Clock/Bmal1*(8, 9) were co-transfected with an hCas9 expression vector, two sgRNA expression vectors targeting *Hes7*, and a plasmid with a puromycin selection marker using FuGENE HD (Promega) as described previously(10). The CRISPR-targeted sequences are as follows: 5’- TGAGAATAGGGACGGCCCCAAGG-3’ and 5’-CAAGGTCTCCAAAACGCGGGCGG-3’. The *Hes7* KO ESC colonies were picked and cultured to establish cell lines after confirmation by sequencing analysis of genomic DNA using primers: 5’-CTTTCCGGGAGCCTCGTGC-3’, 5’-CTTGAGCTGGGCATCTAGGGG-3’, 5’-GACCCACAAATAAAGTTCAGCTCCAAC-3’.

The *Hes7* KO-Rescue cell lines were established by using Tol2 based *Hes7* expression rescue vector, which was constructed by Gibson assembly (New England Biolabs) as follows; the *Hes7* promoter-driven *Achilles* fused *Hes7* gene amplified from pHes7-Achilles-Hes7 (addgene #153528(11)) was inserted between insulator fragments amplified from PB510B-1 (System Biosciences) and assembled into Tol2 based vector with a Zeocin selection marker gene(12). The *Hes7* KO ESCs were co-transfected with the Tol2 based *Hes7* rescue vector and Tol2 transposase expression vector (pCAGGS-TP), and colony picked and cultured to establish *Hes7* KO-rescue cell lines after selection with 15 µg/mL Zeocin.

#### Bioluminescence time lapse imaging and Data analysis of gastruloid

Gastruloids were generated as described previously(13). Briefly, 200 live cells were plated in 40 µL of N2B27 medium into 96-well U bottom plate (Greinier 650185). After 48 h of culture, 150 µL of N2B27 containing 3 µM CHIR99021 was added to each well. At 72 h, 150 µL of medium in each well was replaced with fresh N2B27. At 96 h, the gastruloids embedded in 10% Matrigel (Corning 356231) containing 1 mM luciferin were imaged with a 5-min time resolution under 5% CO_2_ using LV200 system (Olympus). The image sequences were analyzed by ImageJ. Kymographs of the averaged bioluminescence intensity along the segmented lines with 5-pixel width were generated by using the plug-in KymoResliceWide.

#### Quantitative RT-PCR (qPCR)

Gastruloids, ESCs, and *in vitro* differentiated ESCs for 21 days as described previously(12) were washed using ice-cold PBS and total RNA was extracted using Isogen reagent (Nippon Gene) according to the manufacturer’s instructions. First-strand cDNAs were synthesised with 400 or 1,000 ng total RNA using M-MLV reverse transcriptase (Invitrogen) according to the manufacturer’s instructions. Quantitative RT-PCR was performed with iTaq™ Universal SYBR Green Supermix (Bio-Rad Laboratories) by using a StepOnePlus™ Real-Time PCR system (Applied Biosystems). Standard PCR amplification protocols were used and the specificity was confirmed by dissociation-curve analysis. The transcription levels were normalised to the expression level of β-actin. The following primers were used: *Per1*, 5’-CCCAGCTTTACCTGCAGAAG-3’, 5’-ATGGTCGAAAGGAAGCCTCT-3’; *Nr3c1*, 5’-CCTGTCCAGCATGCCGCTATCGAAAA-3’, 5’-CCTGCAGTGGCTTGCTGAATTCCTT-3’; *Bmal1*, 5’-CCACCTCAGAGCCATTGATACA-3’, 5’-GAGCAGGTTTAGTTCCACTTTGTCT-3’; *Clock*, 5’-ATTTCAGCGTTCCCATTTGA-3’, 5’-TGCCAACAAATTTACCTCCAG-3’; *Hes7,* 5’-GAGAGGACCAGGGACCAGA-3’, 5’-TTCGCTCCCTCAAGTAGCC-3’; *Actb*, 5’-GGCTGTATTCCCCTCCATCG-3’, 5’-CCAGTTGGTAACAATGCCATGT-3’.

#### Statistical analysis

Data were analyzed by the statistical methods stated in each legend using GraphPad Prism version 10.0 software or R. Genotype frequencies were compared with expected Mendelian ratios using Pearson’s chi-square goodness-of-fit test. Data normality was assessed with the Shapiro-Wilk test. For two-group comparisons, Welch’s *t* test (parametric) or Wilcoxon rank-sum test (non-parametric) was applied as appropriate. Circadian rhythmicity of corticosterone (24-h period) was assessed with a cosinor regression model(14) using the R package cosinor2(15) v0.4 (function ‘cosinor.lm’, period = 24 h). Rhythmicity of clock gene expression was evaluated using the meta2d function of the MetaCycle R package(16). The main effects of embryonic genotype (WT vs KO), maternal treatment (control vs stress), and their interaction were evaluated by two-way Welch–James heteroscedastic ANOVA using welchADF (v 0.4.2)(17). Approximate F statistics with Satterthwaite-adjusted denominator degrees of freedom (ν) are reported. Pairwise post-hoc comparisons were performed with the Games–Howell test procedure(18). Unless otherwise specified, results with P < 0.05 were considered statistically significant. Data are presented as mean ± SD.

### Supplementary Figures

**Fig. S1.**
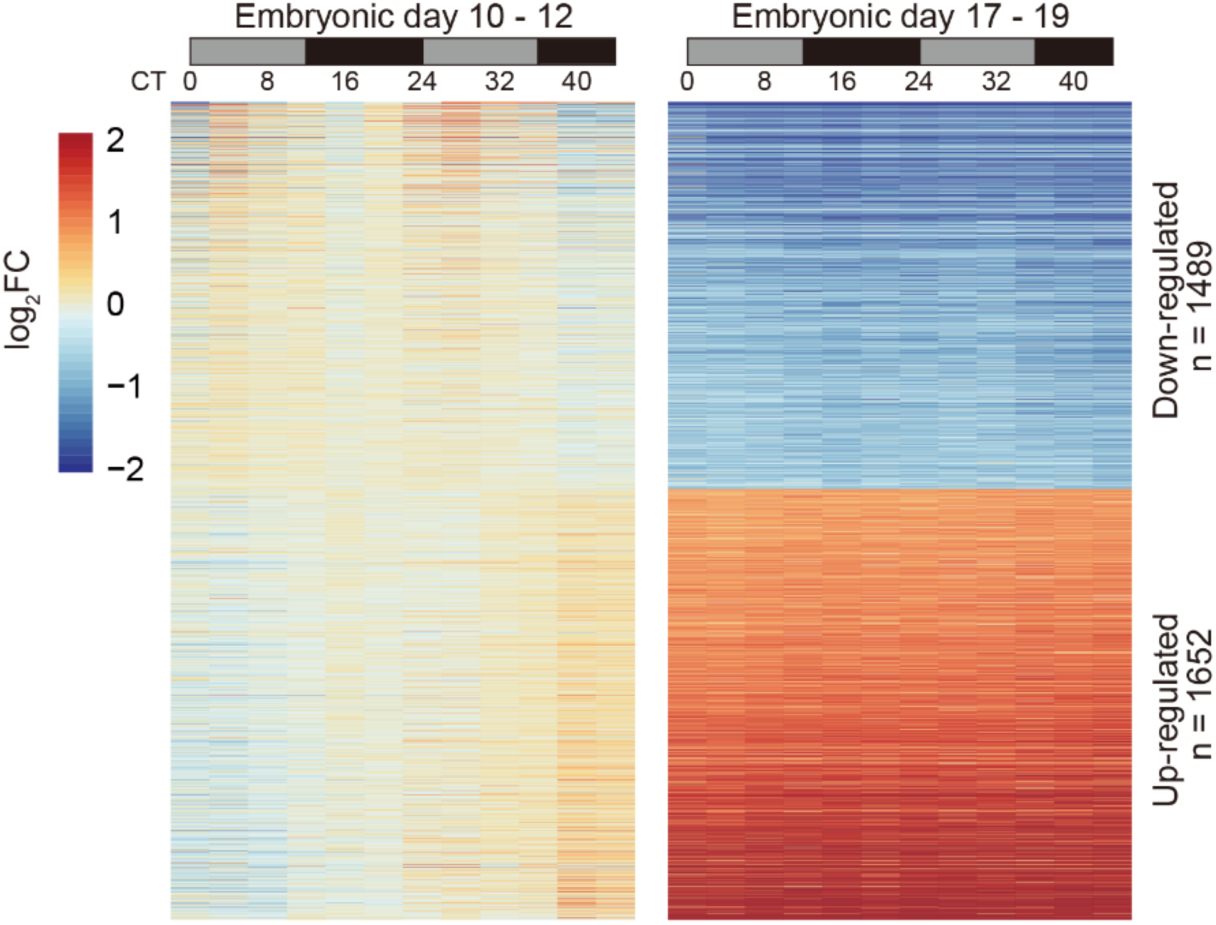
Heatmap view of genes differentially expressed between E10–E12 and E17–E19 mouse embryonic hearts (reanalyzed RNA-seq dataset; DRA005754). Genes with absolute fold change > 2 and FDR < 0.05 are shown (Dataset S1). Rows represent genes (down-regulated, n = 1,489; up-regulated, n = 1,652), and columns represent samples collected at the indicated circadian times (CT0–CT44) on the indicated embryonic days. The gray and black boxes denote the subjective day and night, respectively.

**Fig. S2.**
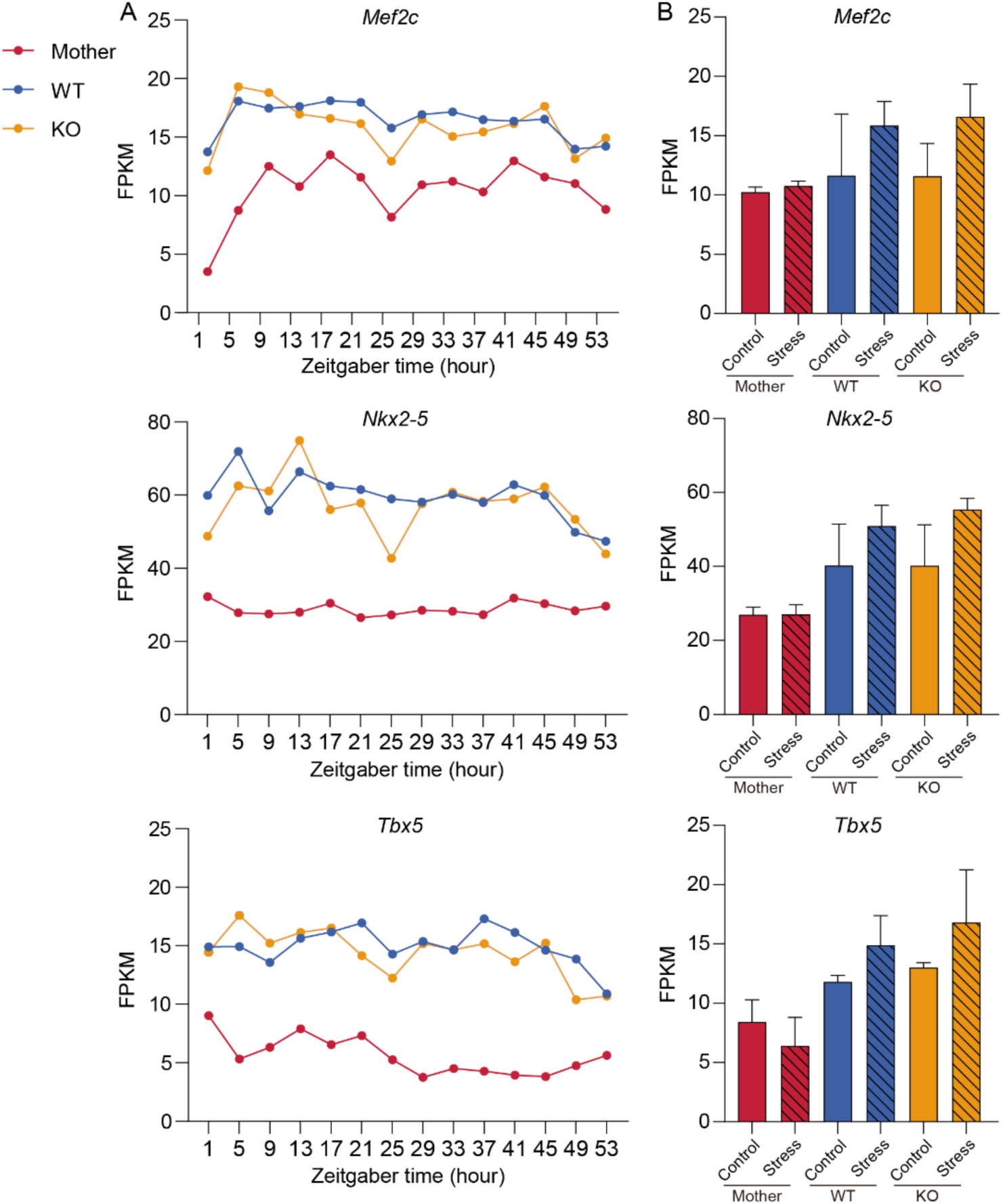
Expression of cardiomyocyte marker genes of maternal and embryonic hearts. Gene expression levels of *Mef2c* (upper), *Nkx2–5* (middle), and *Tbx5* (lower). (A) Time-series expression profiles in hearts sampled every 4 h from ZT1–ZT53 (E10–E12). (B) Maternal and embryonic hearts in the acute-stress experiment. Bars show mean ± SD (n = 3–4 biological replicates).

**Fig. S3.**
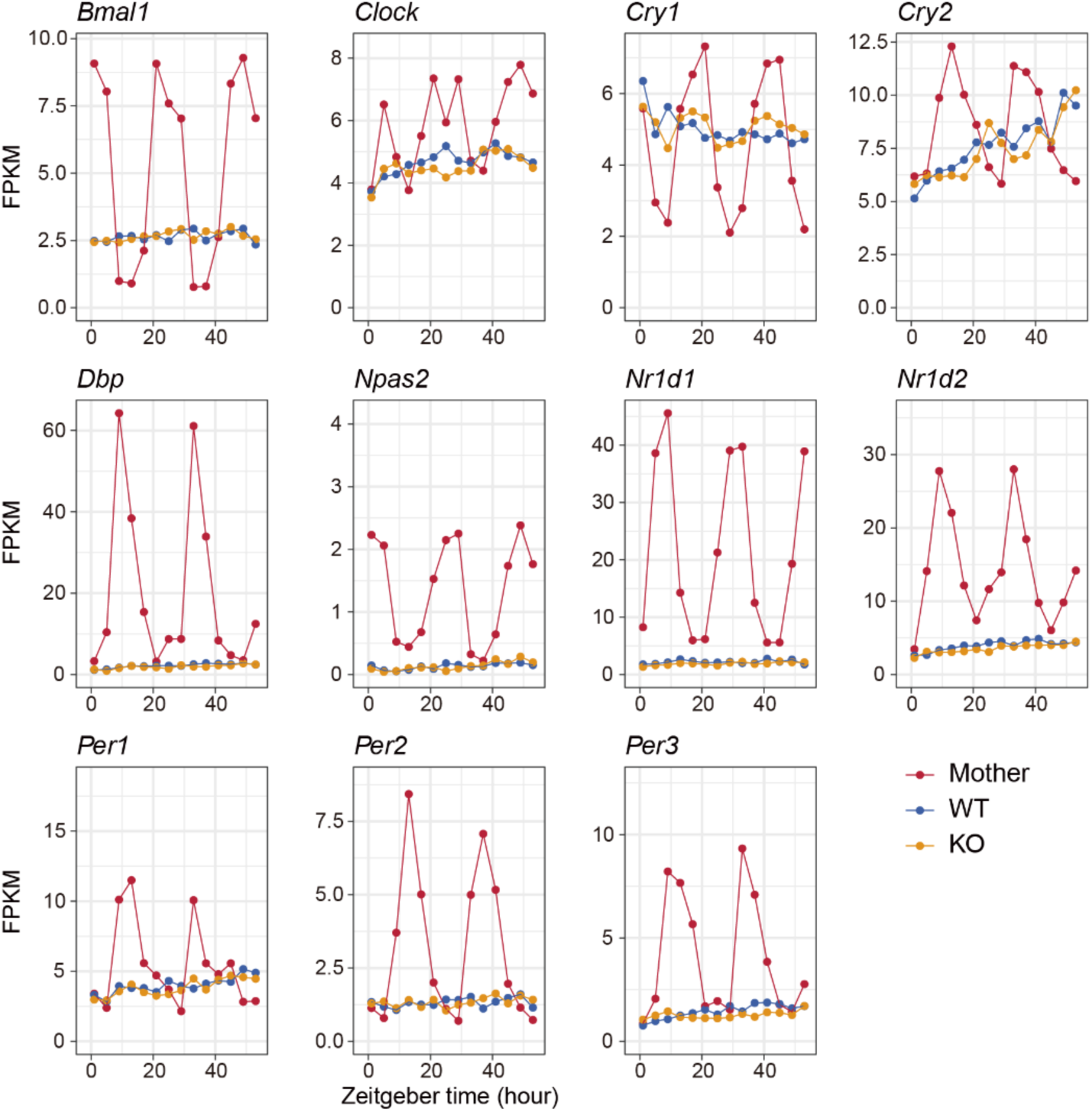
Time-series expression profiles of the indicated clock genes in maternal hearts (red) and WT (blue) or *Hsd11b2* KO (orange) embryonic hearts sampled every 4 h from ZT1–ZT53 (E10–E12). Rhythmicity was assessed by MetaCycle (P < 0.05, relative amplitude > 0.2): all genes except *Clock* were rhythmic in maternal hearts, whereas none were rhythmic in embryonic hearts of either genotype. Embryonic samples were pooled (2–4 hearts per genotype per litter per time point).

**Fig. S4.**
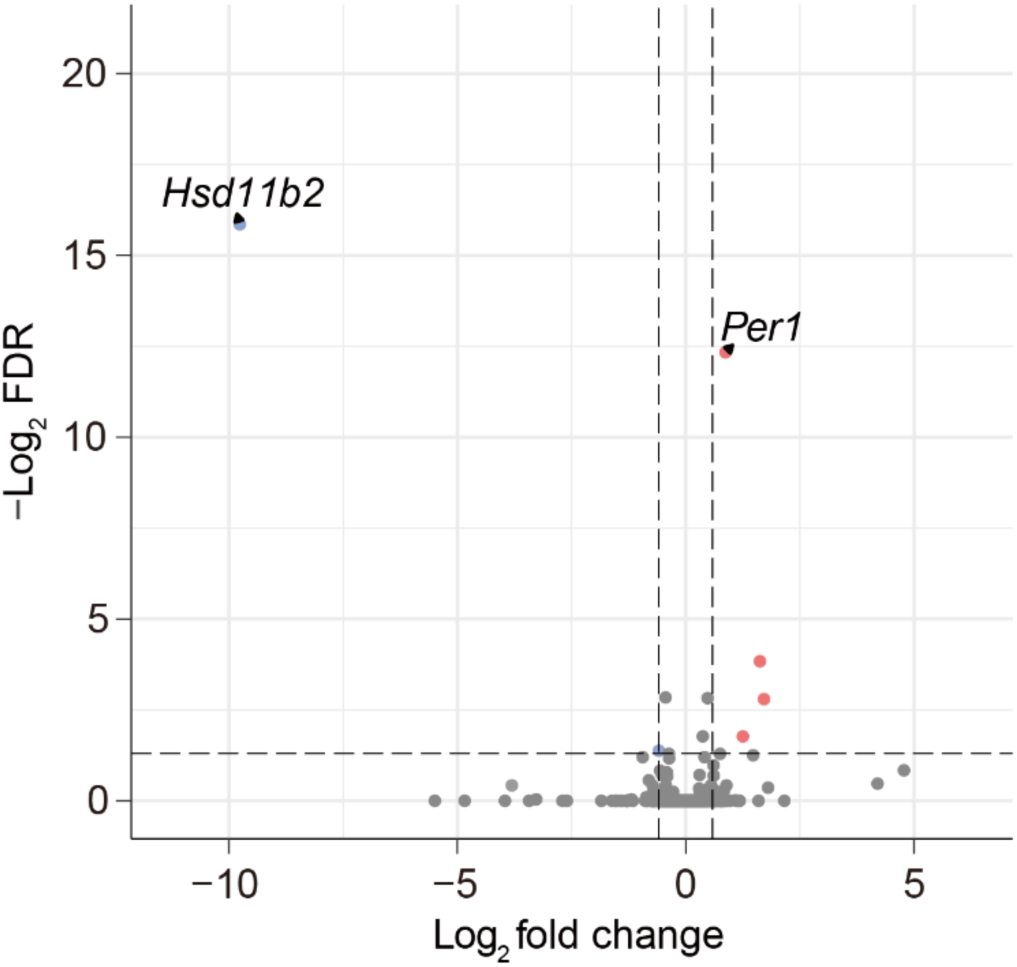
Volcano plot of differentially expression genes comparing WT and *Hsd11b2* KO embryonic hearts collected at E11.5 immediately after a 2-h maternal restraint stress (ended at ZT1). The x-axis indicates log₂ fold change (KO/WT) and the y-axis indicates −log₁₀(FDR). The horizontal dashed line marks FDR = 0.05 and vertical dashed lines denote |log₂FC| = 0.585 (fold change = 1.5). Differentially expressed genes were identified using DESeq2 with thresholds of FDR < 0.05 and absolute fold change > 1.5.

**Fig. S5.**
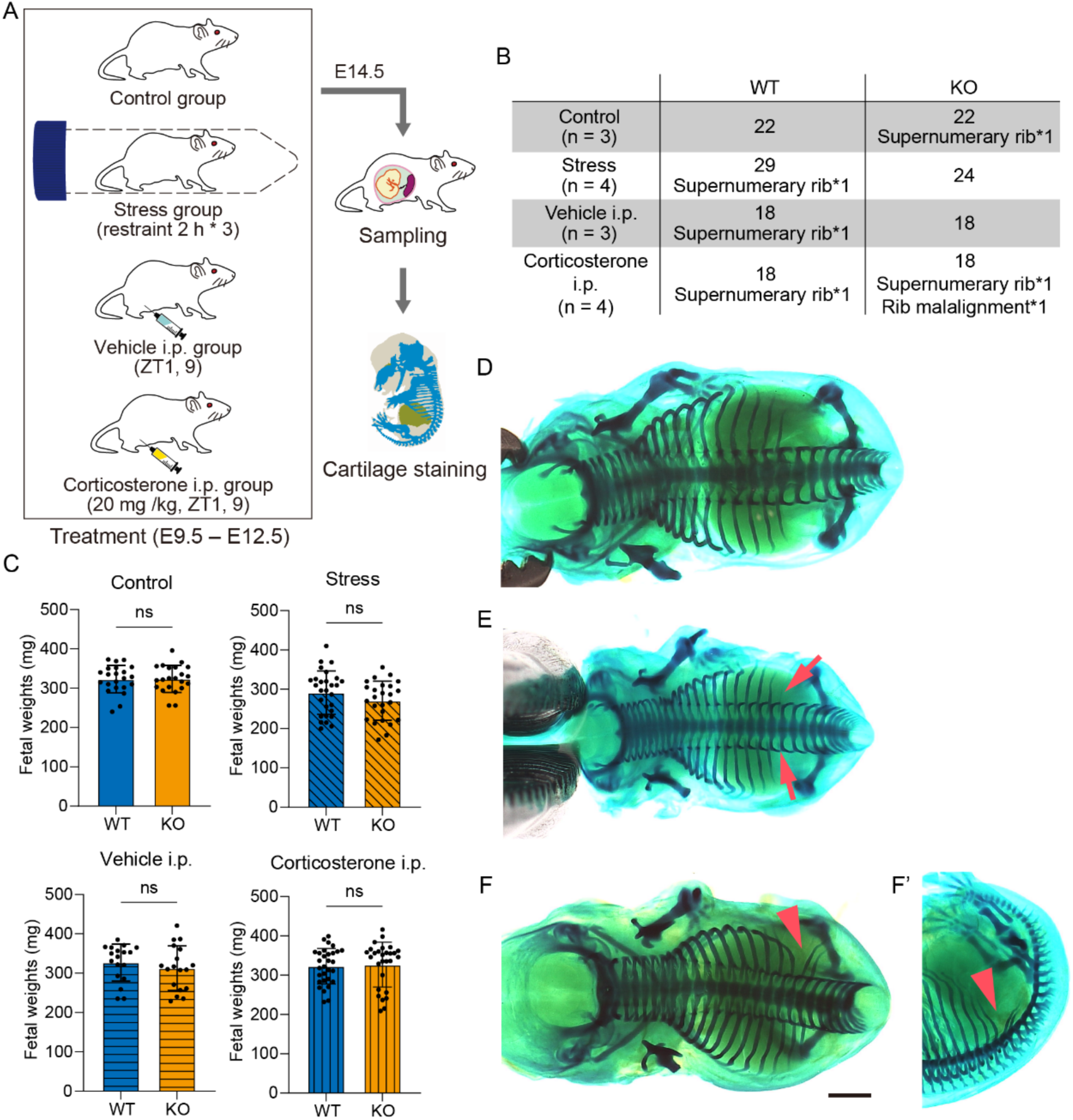
Evaluation of embryonic phenotypes after maternal stress exposure and corticosterone administration. (A) Experimental scheme. Dams carrying co-transferred WT and *Hsd11b2* KO embryos were assigned to four groups and subjected to distinct interventions from E9.5 to E12.5. Embryos were collected at E14.5, and axial skeletal patterning was evaluated by cartilage staining. (B) Table summarizing the numbers of embryos obtained and abnormalities observed in each group. (C) Fetal body weights at E14.5. Bars represent mean ± SD (n, see (B)). ns indicates not significant. (D–F) Dorsal views of E14.5 mouse embryos stained with Alcian blue. (D) Representative normal embryo (Control group, WT). (E) Representative embryo with a supernumerary rib (arrows) (Stress group, WT). (F) Embryo in which the left ribs are deviated caudally (arrowhead) (Corticosterone i.p. group, KO). (F′) Left lateral view of (F). The scale bar represents 1 mm.

**Fig. S6.**
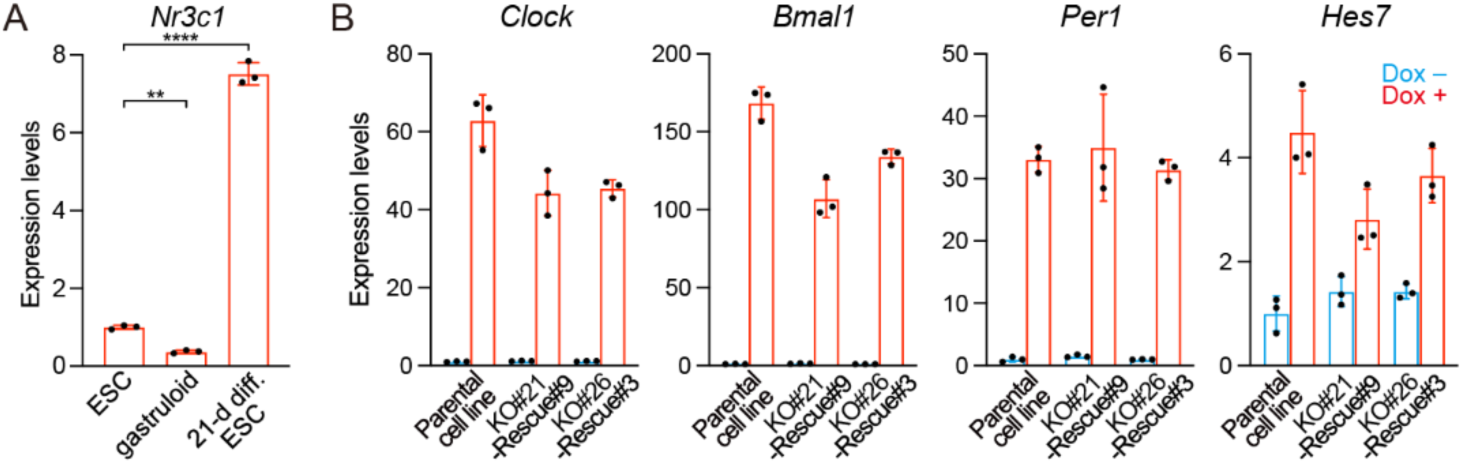
qPCR of *Nr3c1*, *Clock*, *Bmal1*, *Per1,* and *Hes7* mRNA in the indicated cells. (A) *GR* (*Nr3c1*) mRNA in the ESCs, gastruloids, and *in vitro* 21-day differentiated ESCs with circadian oscillations. The average expression level in the ESCs was set to 1. One-way ANOVA followed by a Bonferroni post-hoc test. **P < 0.01, ****P < 0.0001. (B) Colored boxes indicate 1000 ng/mL dox treatment for 2 h (red) or no treatment (blue). The average expression level in the parental cell lines without dox was set to 1. Mean ± SD (n = 3 biological replicates).

**Fig. S7.**
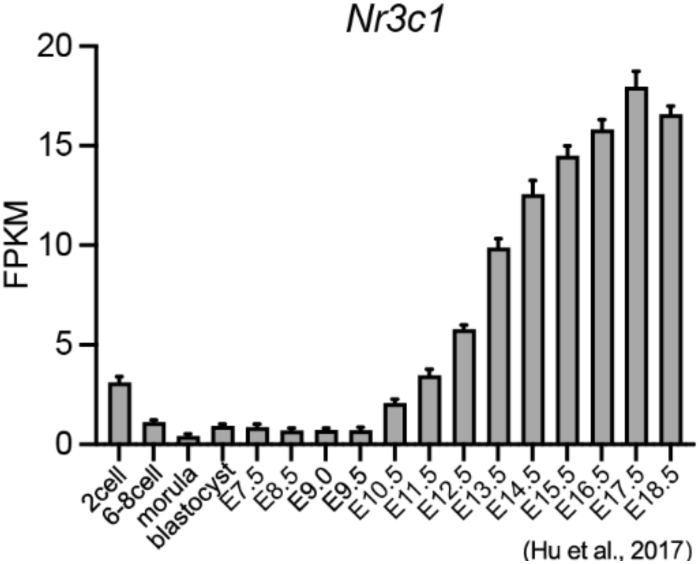
Developmental changes in *Nr3c1* expression. *GR* (*Nr3c1*) mRNA expression at different developmental stages by RNA-seq analysis previously reported(19). Mean ± SD (n = 2-3 biological replicates).

**Dataset S1 (separate file).** List of genes that underwent significant changes in the embryonic hearts from E10–12 to E17–19.

**Dataset S2 (separate file).** List of DEGs in control vs stress conditions in the maternal hearts.

**Dataset S3 (separate file).** The KEGG pathway analyses of the DEGs in control vs stress conditions in the KO embryonic hearts.

